# Endocrine modulation of cortical and retinal blood flow across the menstrual cycle

**DOI:** 10.1101/2024.12.19.629235

**Authors:** Melissa E. Wright, Cassandra Crofts, Saajan Davies, Hannah Chandler, Ian Driver, Michael Germuska, Ylenia Giarratano, Darwin Rashid, Miguel O. Bernabeu, Louise Terry, Jessica J. Steventon, Kevin Murphy

**Affiliations:** School of Physics and Astronomy, Cardiff University Brain Research Imaging Centre (CUBRIC), Cardiff University; Centre for Medical Informatics, Usher Institute, The University of Edinburgh; School of Optometry and Vision Sciences, Cardiff University; School of Medicine, CUBRIC, Cardiff University; Department of Radiology, University of California Davis Medical Center, Sacramento, CA, United States of America

## Abstract

The ovarian hormones, oestrogen and progesterone, have vaso- and neuroprotective effects, likely due to interactions with the cerebrovascular system. This study investigates their neuroendocrine influence on a range of cerebral and retinal vascular functions across a healthy menstrual cycle.

Twenty-six healthy, menstruating females completed imaging sessions and assessment of circulating hormone levels during their early follicular, late follicular, and mid-luteal phase (1-4, 10-12 and 20-22 days after menses onset). Cerebral blood flow (CBF), arterial arrival time (AAT), global oxygen extraction fraction (OEF), cerebrovascular metabolic rate of oxygen (CMRO_2_), carotid artery radius and carotid pulsatility index (PI) were measured using 3T MRI. Retinal vessel density and blood flow resistance were assessed with optical coherence tomography angiography (OCT-A).

Assessed with linear models, increased oestradiol was related to increased global CBF (*χ*^2^(1)=41.682; p=1.074×10^-10^). An independent progesterone increase was also associated with increased global CBF (*χ*^2^(1)=14.979; p=0.0001). In the retina, a relationship was found between oestradiol and decreased retinal blood flow resistance (*χ*^2^(1)=5.28; p=0.0215), which was primarily driven by centrally localised vessels.

This study finds that circulating oestrogen increases blood flow in the eye and brain, while progesterone significantly impacts the brain alone. These effects suggest a potential pathway for neuroprotective mechanisms.

## Introduction

Ovarian hormones, such as oestrogen and progesterone, are considered to have vaso- and neuro-protective effects. A longer reproductive lifespan, and therefore a greater lifetime exposure to oestrogen, is associated with lower cardiovascular disease and dementia risk (1–4), greater cortical volume (5,6), and greater later-life cognitive performance (7). Events associated with a decline in oestrogen levels (e.g., menopause) demonstrate an increased risk of vascular pathology, dementia and glaucoma (8–13). Progesterone is also shown to be protective against progression in a range of animal disease models (e.g., dementia, multiple sclerosis; for a review, see 14). Irregular menstrual cycles show an association with increased cerebrovascular risk (15). Considering the vasodilatory influence of oestrogen reported in animal models (16,17), endocrine modulation of the cerebrovascular system may contribute to this protective effect, especially as oestrogen receptors are found throughout the brain and retinal layers (18–22). Furthermore, modulation of the vascular system is observed following introduction of external hormone manipulation (e.g., hormonal contraceptives [23] or gender-affirming hormone replacement therapy [24,25]). However, the influence of endocrine states on the human cerebrovascular system remains poorly understood.

The menstrual cycle provides an ideal model to investigate the influence of endogenous ovarian hormones on the cerebrovasculature. Across a menstrual cycle, levels of oestrogen and progesterone demonstrate relatively predictable fluctuations that can be utilised to investigate how acute high and low hormone states impact cerebrovasculature. Menstrual hormones have been found to modulate cortical volume (particularly medial temporal structures; 26–31), estimates of brain age (32), neurotransmitter levels (33–35), and resting/task-related cortical function (36–41). These cortical changes may also contribute to menstrual symptoms (e.g., 42–44). To place these results in context and to understand the interactions between ovarian hormones and the neural system, it is important to fully profile cerebrovascular function across this time period.

Early research using transcranial Doppler ultrasound suggests that blood flow metrics differ in the luteal phase, in which both oestrogen and progesterone are elevated (45). Otomo *et al*., (46) investigated differences in quantitative cerebral blood flow (CBF) using pseudo-continuous arterial spin labelling (pCASL) MRI methods and a sample of eight menstruating female participants. They reported an increase in frontal pole CBF in the follicular phase compared to the luteal phase; however, without serum hormone measurements or more detailed information about test days, it is difficult to make conclusions about the specific neuroendocrine mechanisms. Cote *et al.*, (47) also investigated the endocrine modulation of CBF across a menstrual cycle and found oestrogen and progesterone demonstrated opposing and regionally distinct influences. These studies begin to illustrate the impact of menstrual-related hormones on brain vasculature.

The retina represents a more easily imaged section of the central nervous system than the brain, and structural changes have been identified as independent predictors for a range of systemic diseases (48). Optical Coherence Tomography Angiography (OCT-A), which allows for the visualisation of the vessels that permeate and nourish the retina, is highly complementary to vascular MRI. OCT-A has been utilised to demonstrate vessel density changes across a menstrual cycle, particularly the ovulatory phase (49). However, circulating hormone metrics were not collected to allow for mechanistic inference. Recent OCT-A analysis frameworks extract additional informative vascular metrics, including blood flow resistance, to enable a more comprehensive characterisation of retinal vessel properties (50,51). Overall, the literature illustrates short-term, menstrual-related changes on the cerebrovascular system, but this has yet to be investigated with a detailed cerebrovascular battery across brain and eye.

The current study thoroughly catalogues how a range of baseline cerebral and retinal vascular functions vary with menstrual-related endocrine fluctuations in oestradiol and progesterone. Oestradiol is the most potent form of oestrogen and the form most commonly found in reproductive age women, showing regular fluctuations across a menstrual cycle (52). Such a thorough investigation is vital to gain a full picture of the influence of hormones on vascular health, place previous results in better context, and highlight possible mechanisms behind menstrual-related symptoms, such as migraine.

Participants were tested at three time points; the early follicular phase (EFP; where oestradiol and progesterone are at ‘baseline’), the late follicular phase (LFP; where progesterone should remain low but oestrogen peaks), and the mid-luteal phase (MLP; where both hormones are expected to be elevated). As an exploratory aim, we investigated how vascular functions across brain and eye relate to each other and how this coupling differs across high and low endocrine states.

This study therefore has the following aims:

1. Determine the contribution of circulating oestradiol and/or progesterone to variance in brain and retinal baseline vascular function measures across a healthy menstrual cycle.
2. Determine if the relationships between the outcome vascular function measures differ between high and low endocrine conditions.

## Methods

Participants were tested three times across a menstrual cycle, according to the self-reported onset day of menses. The three testing periods were the early follicular phase (EFP; scanned within days 1-4 of a cycle), the late follicular phase (LFP; 10-12 days), and the mid-luteal phase (MLP; 20-22 days). Participants fasted for a minimum of 4 hours before testing and all sessions took place at a similar time of day. Participants were also asked to abstain from caffeine, alcohol, and strenuous physical activity 12 hours before testing sessions, to avoid any influence these may have on haemodynamics (e.g., 53). Each testing session comprised of blood sampling, an MRI scan, and an ocular imaging session.

A summary of outcome variables is presented in Table 1.

**Table 1.** All vascular outcome variables. MRI=Magnetic resonance imaging; MPDL- pCASL=multi post labelling delay pseudocontinuous arterial spin labelling; TRUST=T_2_- relaxation-under-spin-tagging; DEXI-pCASL=dual-excitation pseudocontinuous arterial spin labelling; TOF=Time of flight; DIMAC=dynamic inflow magnitude contrast; OCT-A=Optical Coherence Tomography Angiography.

| Outcome variable | Methodology | Region of interest |
| --- | --- | --- |
| **Cerebral blood flow (CBF)** | MRI – MPLD-pCASL | Desikan-Killiany Atlas (41 regions per hemisphere) |
| **Arterial arrival time (AAT)** | MRI – MPLD-pCASL | Desikan-Killiany Atlas (41 regions per hemisphere) |
| **Oxygen extraction fraction (OEF)** | MRI – TRUST | Global (via sagittal sinus) |
| **Global cerebral metabolic rate of oxygen (CMRO<sub>2</sub>)** | MRI – TRUST and MPLD-pCASL | Global (via sagittal sinus) |
| **Carotid radius** | MRI – TOF | Carotid arteries (left and right) |
| **Pulsatility index (PI)** | MRI – DIMAC | Carotid arteries (left and right) |
| **Retinal vessel density** | OCT-A | Central fovea, parafovea (left and right eyes) |
| **Retinal blood flow resistance** | OCT-A | Central fovea, superior, temporal, nasal, inferior (left and right eyes) |

### Participants

Twenty-six menstruating participants (age mean[SD]=22.98[3.58] years) were recruited. An initial screening session determined eligibility and sought written informed consent. Eligible participants had a regular (i.e., self-reported cycle length between 27-31 days for at least the last three cycles) menstrual cycle. Participants were excluded if they had any commonly accepted contraindications to MRI scanning, a clinically significant condition (such as diabetes or a neurological, cardiovascular, psychiatric, cerebrovascular, or respiratory condition), were pregnant or had been in the last 6 weeks, took hormonal contraceptives presently or in the last 6 months, were post-menopausal, demonstrated an intraocular pressure of >21mmHg (assessed before ocular imaging via an iCare IC100 Tonometer, Mainline Instruments LTD., UK), or had an ocular condition that may impact measurements (e.g., glaucoma or macular disease). Additionally, participants were required to complete a COVID-19 screening questionnaire before each session. Testing sessions were rearranged if participants reported any COVID-related symptoms, and participants were excluded if they were clinically vulnerable or extremely vulnerable to COVID-19. Participants received financial compensation for completed testing sessions. Ethical approval was given by the Cardiff University School of Medicine Research Ethics Committee.

Due to factors such as participant drop out and equipment availability, not all participants contributed to every timepoint and methodology. A breakdown of sample sizes is given in Table 2.

**Table 2.**
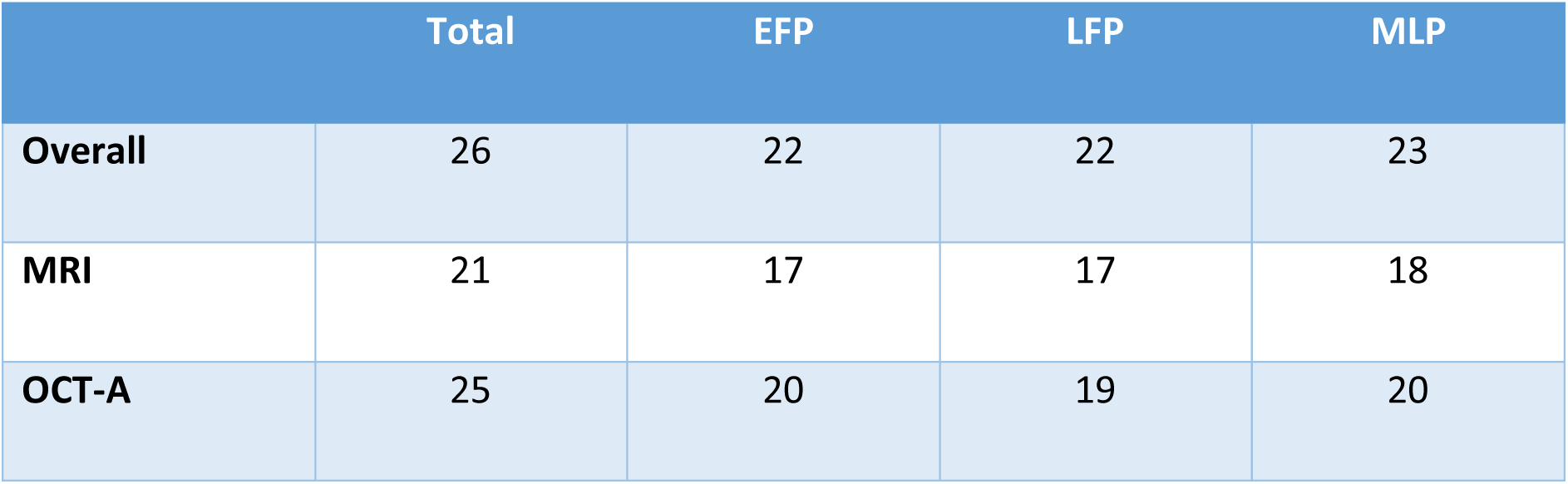
breakdown of participant sample sizes contributing to each timepoint and methodology. EFP=early follicular phase; LFP=late follicular phase; MLP=mid-luteal phase; MRI=magnetic resonance imaging; OCT-A=optical coherence tomography angiography. OCT-A scans were taken in both left and right eyes in each participant.

### Hormone sampling

To assess circulating hormone levels, 5ml of blood was sampled via venepuncture from the hand/arm. The anonymised blood sample was sent to the Medical Biochemistry Laboratories at the University Hospital of Wales for measurement of oestradiol and progesterone serum concentration as NHS standard. Oestradiol levels were measured with an Abbott Architect analyser, while progesterone levels were measured with an Abbott Alinity analyser.

Intercorrelation between these hormones reduces the power of the linear models used; therefore, progesterone values were transformed. Oestradiol levels were regressed from progesterone levels and the model residuals were taken as the variance of progesterone that is independent from oestradiol, referred to as ‘resProgesterone’.

### MRI session – acquisition

Data were acquired at Cardiff University Brain Research Imaging Centre (CUBRIC) on a Siemens MAGNETOM Prisma 3T scanner with a 32-channel head coil. A high-resolution structural T1 image was collected via an MPRAGE scan (1mm^3^; repetition time [TR]=2.1s; echo time [TE]=3.24ms; inversion time [TI]=850ms). The structure of cerebral arteries was imaged using a time-of-flight (TOF) scan (TR=21ms; TE=3.34ms; FA=18; GRAPPA=3; 8 slabs; voxel size=0.6mm^3^).

Global oxygen extraction fraction (OEF) was assessed using a T_2_-relaxation-under-spin-tagging (TRUST; 54,55) sequence (TR=3s; TE=3.9ms, effective TEs=0ms, 40ms, 80ms, 160ms). This was manually positioned over the sagittal sinus, using the AC-PC and occipital notch as navigational anatomical landmarks. A T1 inversion recovery sequence (ΔTR / TE=150ms / 22ms, flip angle=90 degrees, and GRAPPA acceleration factor=2; with 960 acquisitions > 16 repeats of 60 measurements) was also collected to quantify the T1 of venous blood. This was used to calculate haemoglobin (Hb; 56), necessary for estimating OEF and then CMRO_2_ (cerebral metabolic rate of oxygen). An in-house multi post-labelling delay pseudocontinuous arterial spin labelling (MPLD-pCASL) perfusion scan was completed to assess cerebral blood flow (CBF) and arterial arrival time (AAT), with the tagging plane positioned perpendicular to the internal carotid arteries using the TOF scan as reference (maximum TR=5.6s; TE=11s; voxel resolution=3.4x3.4x6.0mm; tag duration=1500; post-labelling delays [PLDs]=250-3000ms in steps of 250ms; GRAPPA=2). Siemens in-built prescan normalize correction was applied to remove spatial signal inhomogeneity due to receive array coil sensitivity. To allow quantification, two separate equilibrium magnetization maps (M0 scan; phase encoding direction PA and AP) were also obtained, with TR=6000ms and TE=11ms.

Pulsatility index (PI) in the internal carotid arteries was measured using dynamic inflow magnitude contrast (DIMAC) imaging (57). A fast single-slice echo planar imaging (EPI) image was acquired, which suppresses static signal leaving the inflowing blood signal in the flow-velocity regime (57,58). This provided flow-velocity-weighted images with a 15ms temporal resolution, allowing the pulse waveform to be resolved for individual heartbeats. The single EPI slice was acquired perpendicular to the internal carotid arteries (TR=15ms; TE=6.8ms; flip angle=90°; 2x2×10mm; 200x200mm field of view; GRAPPA 5; phase partial Fourier 0.75; 4096 repetitions in 65s; slice thickness=10mm).

To monitor and measure physiological data during the scan, a pulse oximeter (Biopac Systems Inc, California, USA), nasal cannula (AD Instruments gas analyser), and custom-built respiratory belt were secured.

### MRI pre-processing

*Cerebral blood flow and arterial arrival time* – Pre-processing of the pCASL sequence was completed using the Analysis of Functional NeuroImages (AFNI) software package (59; provided in the public domain by the National Institutes of Health, Bethesda, MA, USA; http://afni.nimh.nih.gov/afni). Following motion correction, the scan was split into separate PLDs (5 pairs of each) and the tag minus control signal change in magnetisation was calculated for each. For quantification, the M_0_ of blood was calculated as:

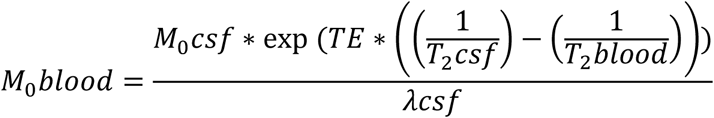

*λcsf* = blood-CSF partition coefficient (1.15; 60,61)

*M*_0_*csf* =the M_0_ of the cerebral spinal fluid (CSF) was taken from a CSF mask generated around a manually positioned central point in the lateral ventricles.

*TE* =echo time (0.019s)

*T*_2_*csf* =T_2_* of CSF (0.4; 60)

*T*_2_*blood* =T_2_* of blood (0.06s; 62)

Using a non-linear fitting algorithm (AFNI’s 3dNLfim function), the general kinetic perfusion model provided final quantification (63). Final CBF and AAT maps were thresholded by a goodness-of-fit measure (R^2^> 0.6) to remove spurious model fits.

FLIRT (64) was used to generate a transformation matrix to register the Desikan-Killiany Atlas to each participants’ native space. The cortical regions were further restricted to the individual’s grey matter voxels (for more information, see Supplemental material). The median CBF and AAT were extracted for each region from voxels that passed threshold (65). The average CBF and AAT across all surviving voxels were taken as global measures for comparison.

*Global oxygen extraction fraction and cerebral metabolic rate of oxygen –* Global OEF and CMRO_2_ metrics were estimated using TRUST scan data and global grey matter CBF values estimated from the PCASL data and a grey matter mask. The T2 of blood was calculated by non-linear least squared fitting of a mono-exponential equation to the TRUST difference data, as a function of the effective echo times and T_2_. The average of the two most intense voxels from a sagittal sinus regions-of-interest (ROI) was used and venous oxygenation (Yv) was estimated by inverting the relationship between Yv, Hb and the T_2_ (54). OEF was calculated as:

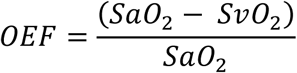

Global CMRO_2_ was calculated as:

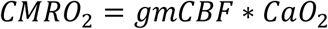

where CaO_2_ is defined as:

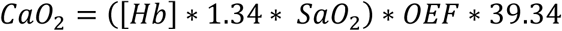

SaO_2_ was assumed to be 98% as this was a healthy young cohort.

*Time-of-Flight analysis* – TOF scans were analysed to obtain the average radius of the carotid arteries. Using the Vascular Modelling Toolkit (VMTK; 66), the ROI, defined as the length of artery between the carotid scan slice position and the first branch of the middle cerebral artery, was extracted and converted into a 3D surface projection. Centrelines were plotted and the average vessel radius was calculated by plotting a Voronoi diagram of the closest points on the surface projection to each point on the centreline. The carotid radius was extracted for the left and right side separately. The surface model was separated into two halves and the carotid arteries in each surface were manually separated from the halves of the Circle of Willis. Non-carotid vessels or vessel fragments were then removed using VMTK’s surface connectivity function, which removes all structures except the largest in the model. When either carotid could not be isolated due to noise, only one artery was included in the analysis (EFP=2, LFP=1, MLP=2). When data for both carotid arteries were not sufficiently high quality to extract the carotid surface, that participant was excluded (EFP=3, LFP=5, MLP=4).

*Carotid pulsatility index* – Left and right internal carotid artery DIMAC timeseries were formed by averaging over a 4x4 voxel square region, centred on each artery. A 3s cut-off high-pass filter and 5^th^-order (21 timepoint frame length) Savitzky-Golay low-pass filter was applied to each timeseries, then each pulse period was modelled by a Fourier series basis set, with 5 sine/cosine pairs with periods ranging from one to one fifth of the pulse period. PI was calculated as the range of the modelled timeseries divided by the mean for each pulse period, then averaged across all pulse periods in the scan.

### OCT session acquisition

OCT-A images were acquired on a Triton Swept-Source OCT device (TOPCON healthcare [Great Britain] medical limited, Newbury, UK). A fovea-centred OCT-A scan was acquired in both eyes (scan area=3mmx3mm).

Participants were instructed to fixate on a central cross and remain as still as possible. At least two images were taken in each eye, with at least a 60/100 quality rating (calculated by the instrument). These two images underwent manual subjective review by author MEW and the best quality scan, as defined by a lack of artefacts such as motion or blurring, was taken forward for further analysis. For one participant (all three phases), no OCT-A scans of sufficient quality were available, so they were excluded from the OCT-A analysis. For one timepoint in one participant, only the left eye provided sufficient image quality (1 MLP).

Eye-tracking was enabled to minimise the contribution of eye movements. All images were obtained with minimal room lighting to ensure maximum natural pupil size.

### OCT-A pre-processing

Vessel density and retinal blood flow resistance were estimated. Initially, a 2D image of the superficial retinal layer (from the internal limiting membrane layer to the inner plexiform layer, as defined by the instrument) was extracted and binarized to produce an image comprised of vessel (white) and non-vessel (black) pixels (see Figure 1).

**Figure 1.**
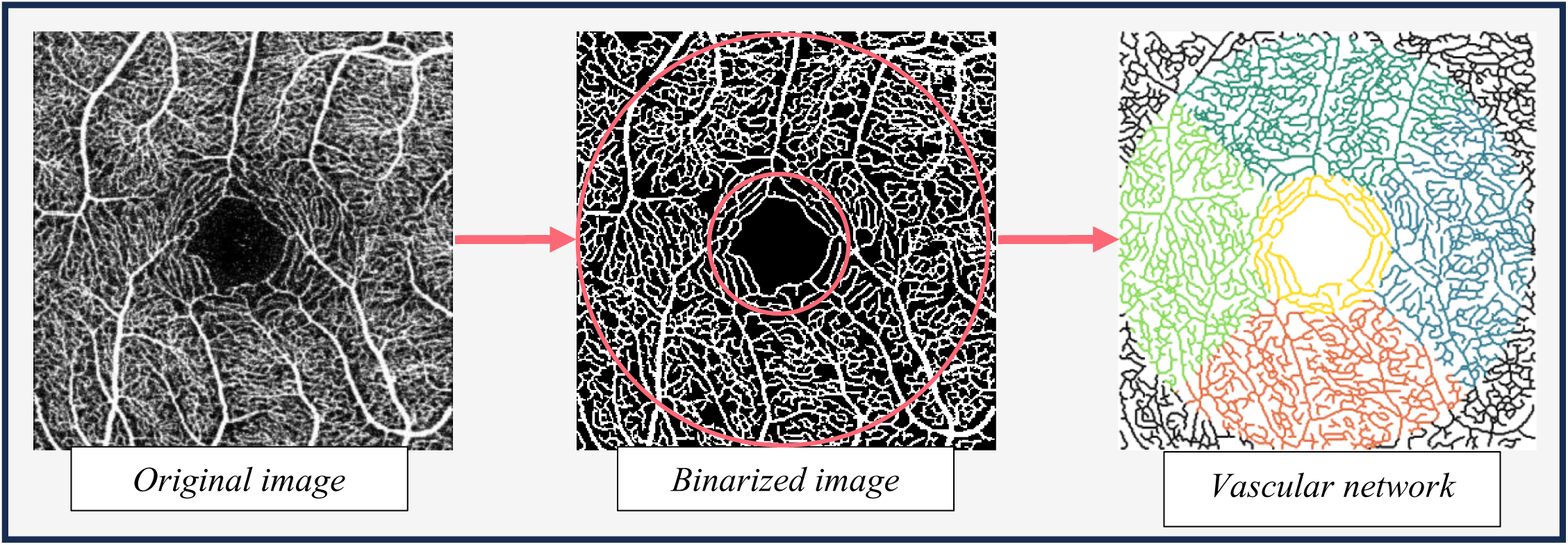
Illustration of the OCT-A analysis pipeline. Red rings =fovea (1mm diameter) and parafovea (3mm diameter) areas used to extract vessel density metrics. Coloured area =fovea, temporal, superior, nasal, and inferior regions used for extracting retinal blood flow resistance metrics.

This binarized image was used as the input to a bespoke analysis software. A detailed explanation of the software has been provided previously (50,51). In brief, the scans undergo automated segmentation using the optimally oriented flux vessel enhancement filter, followed by thresholding (67). Post-processing cleaning was performed to remove segmentation artefacts (i.e., small, isolated clusters of white pixels less than 30 pixels). The vasculature captured in the binarized image was then skeletonised and modelled as a network with nodes representing white pixels along the vessel centreline and edges connecting neighbouring pixels. The network was embedded into a Euclidean space and each node associated with the corresponding coordinates and vessel radius — distance between pixel in the centreline and the closest non-vessel pixel. Features were then extracted from the binary mask and the vascular network. The ratio of vessel pixels to overall number of pixels in the region of interest was extracted as the vessel density in the central fovea and parafovea (see Figure 1), whereas retinal blood flow resistance was calculated using Poiseuille’s law at each vessel segment as follows:

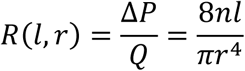

ΔP =pressure difference between two ends of a vascular segment

*Q* =flow rate

*l* =length

*r* =radius

The constant viscosity of blood is assumed to be:

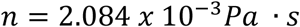

The mean blood flow resistance was extracted for each ROI (the central fovea and superior, temporal, inferior, and nasal regions; see Figure 1) and taken forward for further analysis.

### Statistical Analysis

The contribution of circulating hormone levels to the variance in the outcome variables was calculated using linear mixed models. Statistical analysis was completed in R software (v.4.2.2; 68) and linear models were constructed using the ‘lmerTest’ analysis package (69). Statistical significance was set at the P<0.05 level. With all models, residual plots were visually inspected and showed no notable deviations from homoscedasticity or normality.

Oestradiol, progesterone, and ROI (if applicable) were used as fixed effects, with participant as a random effect. If a statistically significant main effect of hormone was found, an interaction term between ROI and hormone levels was added to investigate if the effect is driven by a particular ROI (if applicable). In all cases where there are more than two ROIs, the average value over all ROIs was included as a global comparison. For OCT-A and carotid metrics, both eyes/vessels were included in the analysis. The laterality (left or right) was inputted as a fixed effect in order to account for similarity between eyes/vessels.

The goal of the second aim was to investigate the relationships between vascular outcomes (detailed above; see Table 1) and investigate how these relationships differ by low or high hormonal state. To address this exploratory aim, the highest and lowest hormone phase for each participant was identified (if two phases had the same level, the earlier phase [i.e., EFP > LFP > MLP] was used for the low hormone condition, and the later phase [i.e., MLP > LFP > EFP] was used for the high hormone condition). High and low groups for oestradiol and progesterone were grouped separately, leading to four conditions. The data were standardised and a cross-correlation matrix generated (Pearson’s rho). A principal components analysis (PCA) was used on each correlation matrix to reduce the dimensionality of the data and investigate shared variance across the variables.

## Results

### Hormones

The levels of circulating oestradiol and progesterone for all participants are presented in Figure 2. The expected LFP oestrogen peak was not captured in all participants and there was a statistically significant correlation between levels of circulating oestradiol and progesterone (all participants and timepoints included; r(65)=0.463; p=7.95×10^-5^).

**Figure 2.**
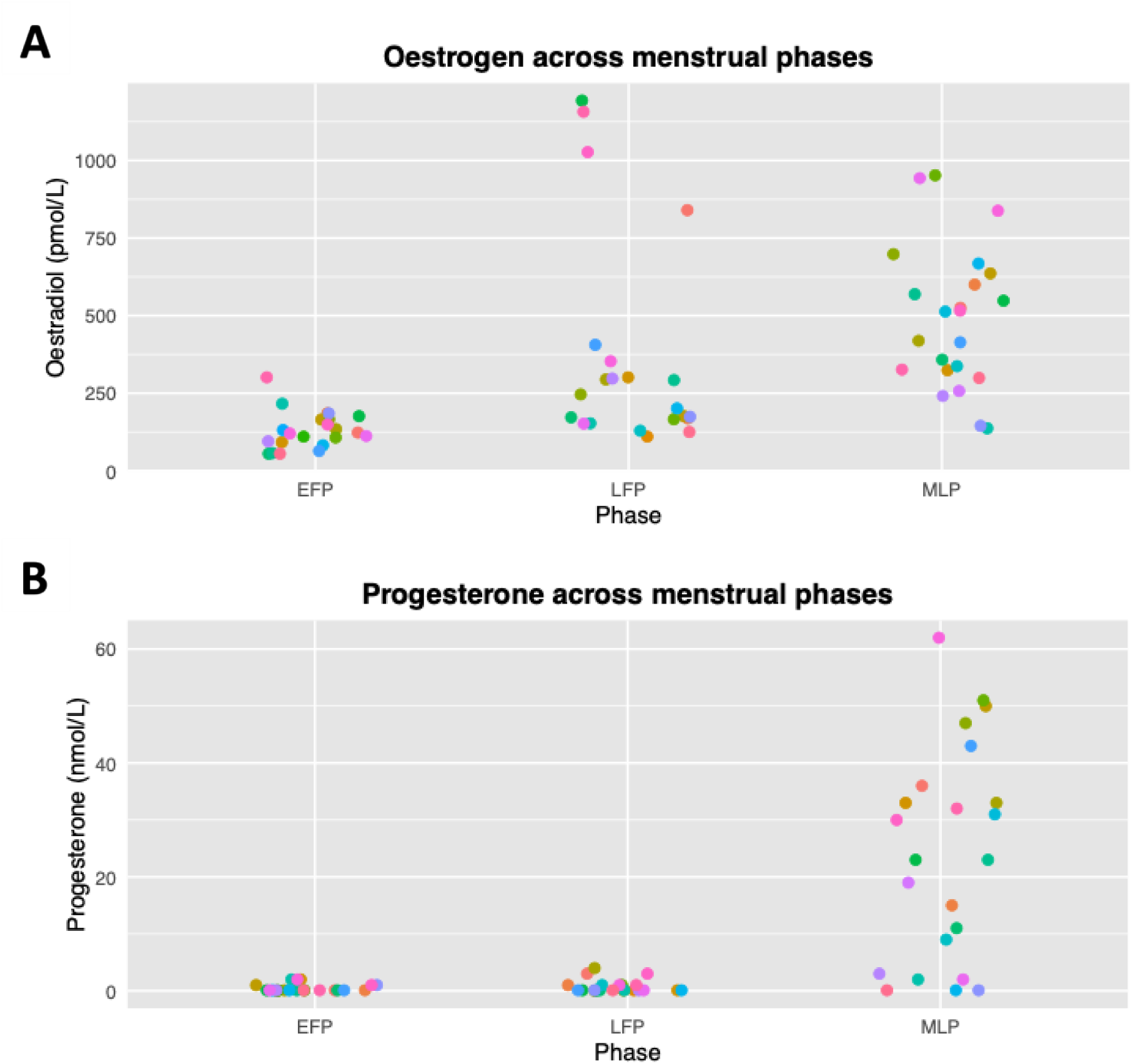
Levels of circulating hormones across the three tested menstrual cycle phases. Panel A =oestrogen (in the form of oestradiol [pmol/L]); panel B =progesterone (nmol/L); EFP =early follicular phase; LFP =late follicular phase; MLP =mid-luteal phase. Coloured by individual participant.

Progesterone values were therefore transformed to avoid issues of intercorrelation. This value is referred to as ‘resProgesterone’.

### Cerebral blood flow

Oestradiol was found to explain a significant amount of global CBF variance (*χ*^2^(1)=41.682; p=1.074×10^-10^), with global CBF increasing by 6.995×10^-3^ml/100g/min ± 1.081×10^-3^ (standard error [SE]) with each unit increase of oestradiol. This represents an average variation of 3.552 ml/100g/min across a menstrual cycle (based on the average max-min of hormone level in participants with more than one timepoint [507pmol/L], though previous literature suggests the fluctuation in oestradiol across a cycle can be up to 2x larger; ,29,70,71). Levels of resProgesterone also explained a significant amount of CBF variance (*χ*^2^(1)=14.979; p=0.0001), with CBF increasing by 0.08087ml/100g/min ± 0.02088 (SE) with each unit increase of resProgesterone, representing a change of 1.652 ml/100g/min across a menstrual cycle (assuming a difference of 20.433nmol/L resProgesterone).

Interaction terms between ROI and each set of hormone levels did not explain a significant amount of variance (i.e., the relationship between the hormone and CBF in each ROI didn’t significantly differ from that globally; oestradiol: *χ*^2^(83)=72.53; p=0.787; resProgesterone: *χ*^2^(83)=96.311; p=0.151).

### Arterial arrival time

Oestradiol did not explain a significant amount of AAT variance (*χ*^2^(1)= 0.0003; p=0.986). Levels of resProgesterone were also not found to explain a significant amount of AAT variance (*χ*^2^(1)= 0.0005; p=0.982).

### Global oxygen extraction fraction

Neither oestradiol (*χ*^2^(1)=0.451; p=0.502) nor resProgesterone (*χ*^2^(1)=0.381; p=0.537) explained a significant amount of global OEF variance.

### Global cerebral metabolic rate of oxygen

As above, neither oestradiol (*χ*^2^(1)=0.303; p=0.582) nor resProgesterone (*χ*^2^(1)=0.546; p=0.460) explained a significant amount of global CMRO_2_ variance.

### Carotid radius

In the carotid radius model, one datapoint with an outlier model residual value (defined as 2.5*SD away from the mean) was removed due violation of the model assumptions (linearity and heteroscedascity; further investigation found this datapoint also had an outlier radius value). Neither oestradiol (*χ*^2^(1)=0.017; p=0.895) nor resProgesterone (*χ*^2^(1)=0.849; p=0.357) explained a significant amount of carotid radius variance.

### Carotid pulsatility index (PI)

Residual plots of the carotid PI model were examined, and two datapoints excluded for outlier residual values that caused assumption violations. After examination of the hormone main effects, neither oestradiol (*χ*^2^(1)=0.758; p=0.860) nor resProgesterone (*χ*^2^(1)=1.029; p=0.794) explained a significant amount of carotid PI variance.

### Retinal vessel density

Neither oestradiol (*χ*^2^(1)=0.002; p=0.964) nor resProgesterone (*χ*^2^(1)=0.216; p=0.642) explained a significant amount of vessel density variance.

### Retinal blood flow resistance

Oestradiol was found to explain a significant amount of resistance variance (*χ*^2^(1)=5.283; p=0.0215), decreasing resistance by -1.868×10^-8^ Pa·s/μm^3^ ± 8.107×10^-9^ (SE) with each unit increase of oestradiol. This translates to a change of blood flow resistance across the menstrual cycle of -1.2×10^-4^ Pa·s/μm^3^. The resProgesterone factor, however, did not significantly contribute to the model (*χ*^2^(1)=0.395; p=0.530).

This interaction between oestradiol and ROI was significant (*χ*^2^(5)=11.603; p=0.0407). By inspecting the individual fixed effects, the oestradiol-resistance relationship was found to be significantly greater in the central (foveal) region compared to the global average (t(6.765×10^2^)=-2.136; p=0.033; estimate=-5.171×10^-8^; SE=2.421×10^-8^), indicating that this region may be driving the result. The other ROIs did not significantly differ from the global average (nasal: t(6.765×10^2^)=0.568, p=0.571; inferior: t(6.765×10^2^)=0.335, p=0.738; temporal=t(6.765×10^2^)=0.832, p=0.406; superior: t(6.765×10^2^)=0.402, p=0.688).

A summary of results is provided in Table 3. Endocrine influences on haemoglobin levels and end-tidal CO_2_ were also examined (see Supplemental material) but no statistically significant effects were found, suggesting this did not influence these results.

**Table 3.** Summary of results from all vascular outcome variables. . n.s. =not significant.

| Outcome | Hormone | $\chi^2$<br>statistic | p-value | Regional<br>effect | Direction |
| --- | --- | --- | --- | --- | --- |
| **CBF** | Oestradiol | 41.682 | <0.001 | n.s. | Increase |
|  | Progesterone | 14.979 | <0.001 | n.s. | Increase |
| **AAT** | Oestradiol | 0.0003 | n.s. | - | = |
|  | Progesterone | 1.604 | n.s. | - | = |
| **OEF** | Oestradiol | 0.451 | n.s. | Global | = |
|  | Progesterone | 0.381 | n.s. | Global | = |
| **CMRO<sup>2</sup>** | Oestradiol | 0.363 | n.s. | Global | = |
|  | Progesterone | 1.061 | n.s. | Global | = |
| **Carotid radius** | Oestradiol | 0.017 | n.s. | - | = |
|  | Progesterone | 0.849 | n.s. | - | = |
| **PI** | Oestradiol | 0.758 | n.s. | - | = |
|  | Progesterone | 1.029 | n.s. | - | = |
| **Retinal vessel**<br>**density** | Oestradiol | 0.002 | n.s. | - | = |
|  | Progesterone | 0.216 | n.s. | - | = |
| **Retinal blood**<br>**flow resistance** | Oestradiol | 5.283 | <0.05 | Yes | Decrease |
|  | Progesterone | 0.395 | n.s. | - | = |

### Exploratory relationships analysis

In an exploratory analysis, we investigated how the relationships between vascular functions differ in high and low endocrine state. While this study was not designed to test this research question, this data can give an idea of whether there is a pattern in how low/high hormone states impact vascular functions relationships, which can then be specifically investigated in the future.

PCA was completed on all conditions (low oestrogen, high oestrogen, low progesterone, high progesterone; cross-correlation tables and scree plots available in the supplemental material). In all conditions, the first two components explained at least 70% of the variance and the loadings of all variables onto these components were plotted. As can be seen from figure 3, the variables are notably more clustered in the low hormone condition for oestrogen and progesterone than the high hormone condition, suggested that there is less shared variance in the high endocrine conditions. The data from the high hormone conditions were then projected to the principal component space generated by the corresponding low hormone condition. Again, the first two components were examined.

**Figure 3.**
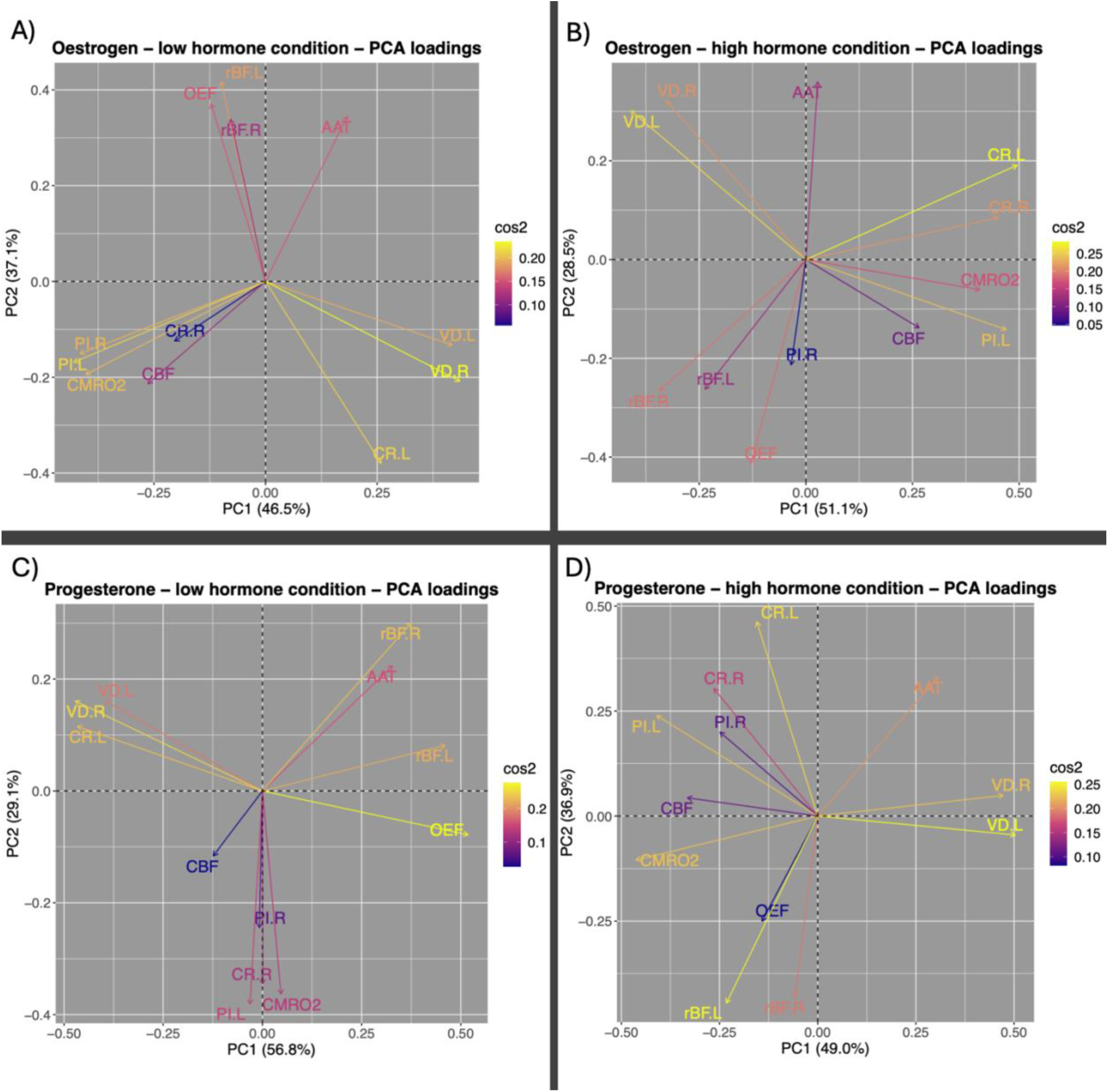
Loadings of all variables on the two extracted principal components (PC1 and PC2) following principal component analysis (PCA) of each condition (low oestrogen, high oestrogen, low progesterone, high progesterone). Cos2 refers to the importance of that component/dimension to that particular variable. AAT=Arterial arrival time; CBF=Cerebral blood flow; CR.L=Carotid radius left; CR.R=Carotid radius right; CMRO_2_=Cerebral metabolic rate of oxygen; OEF=Oxygen extraction fraction; PI.L=Pulsatility index left; PI.R=Pulsatility index right; rBF.L=Retinal blood flow resistance left; rBF.R=Retinal blood flow resistance right; VD.L=Retinal vessel density left; VD.L=Retinal vessel density left.

The amount to which each participant’s data aligned with each component was reduced in the high hormone condition by a small percentage in 3/4 components (Oestrogen component 1: -1.357%; oestradiol component 2: -1.809%; progesterone component 1: -14.406%; progesterone component 2: +3.294%; data plotted in figure 4). For the low oestrogen condition, there was roughly a greater contribution of most of the cortical metrics and retinal vessel density in the first component, while the second component had greater loading from retinal blood flow, AAT, and OEF. For the low progesterone components, roughly the retinal metrics, OEF, and AAT showed greater contributions to the first component, while the others demonstrated greater loadings to the second.

**Figure 4.**
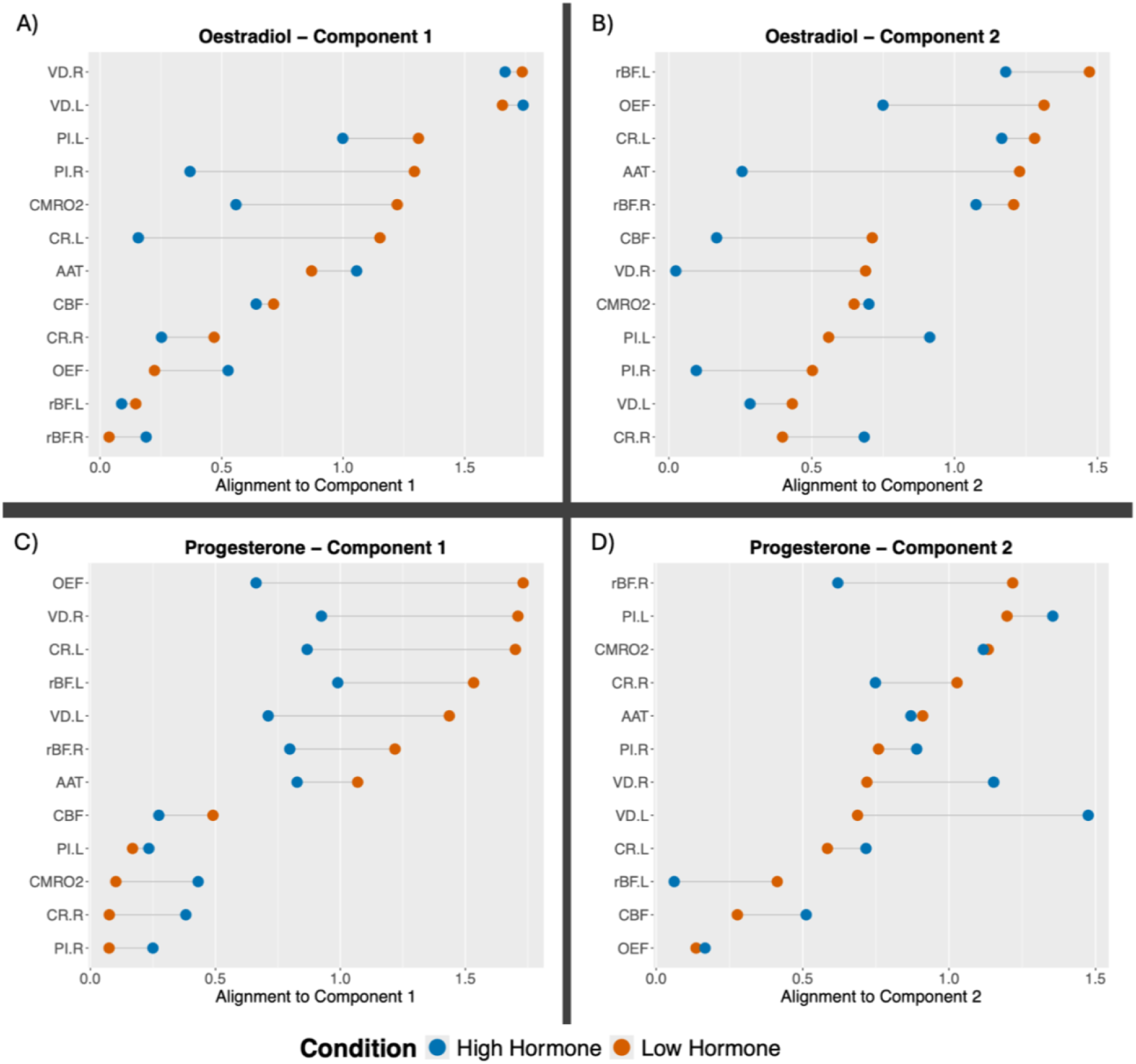
The amount to which each vascular outcome variable (from either the high or low hormone condition) is aligned with each principal component plotted for the oestradiol and progesterone conditions. Blue datapoints are from the high hormone condition (either oestradiol [A and B] or progesterone [C and D]), while orange datapoints are from the low hormone condition (either oestradiol [A and B] or progesterone [C and D]). AAT=Arterial arrival time; CBF=Cerebral blood flow; CR.L=Carotid radius left; CR.R=Carotid radius right; CMRO_2_=Cerebral metabolic rate of oxygen; OEF=Oxygen extraction fraction; PI.L=Pulsatility index left; PI.R=Pulsatility index right; rBF.L=Retinal blood flow resistance left; rBF.R=Retinal blood flow resistance right; VD.L=Retinal vessel density left; VD.L=Retinal vessel density left.

## Discussion

This study investigated the neuroendocrine influence of menstrual hormones (i.e., oestrogen and progesterone) on a battery of baseline cerebrovascular and retinal vascular functions. Animal research has suggested that oestrogen increases vasodilation and CBF (16,17), maintains blood-brain barrier function (72), promotes angiogenesis (17), and improves recovery processes after ischemic injury (73,74), all of which are longer term processes. Progesterone has also been linked to brain injury recovery (75–77), vascular reactivity (78,79) and inflammation (though evidence for the direction of this effect differs; ,80,81). Early work has started to find variations in baseline vascular functions sensitive to menstrual-related endocrine fluctuations (45–47,49), but this has yet to be investigated with a thorough cerebrovascular battery across both brain and eye. This is important to establish as changes in cerebrovascular function will in turn influence the health of the surrounding neural tissue and will also impact more dynamic vascular functions, such as vascular reactivity and the hemodynamic response function.

Oestradiol and additional progesterone variance were modelled so their individual contributions could be calculated. We found that oestradiol regressor was significantly associated with increased CBF, which is in line with previous animal and human research (e.g., 16,47). Progesterone also significantly contributed to increasing CBF. While the CBF change is only ∼8% across the menstrual cycle, it is notable when considering it occurs over the regular fluctuations of the menstrual cycle, which are relatively small hormonal changes compared to other key life changes, such as pregnancy or menopause. Contrary to previous data by Cote *et al.*, (47), we found that these neuroendocrine effects occurred globally across the cortex.

The influence of hormones on baseline CBF has a number of implications. Firstly, blood oxygen level dependant (BOLD) fMRI is a common technique for investigating neural activation. The BOLD signal is a complex phenomenon that reflects both neural activity and vascular reactivity, with baseline vascular function impacting the final BOLD response (82). Endocrine-dependant changes in CBF may therefore lead to extra noise in studies with menstruating participants, especially if comparing groups or timepoints associated with different endocrine states (e.g., menstrual cycle phase, menopause). Measuring hormone levels would be an ideal way of accounting for and controlling for this factor, but controls could also be more easily implemented by asking the time since last period to account for menstrual phase or by testing during the EFP (when hormones are expected to be at ‘baseline’). As CBF changes can affect volumetric measurements, these results are also important to place menstrual-related investigations of cortical structure into better context. For example, as our observed neuroendocrine effects were global, it provides more confidence that regionally-specific menstrual changes to volume (e.g., 26,28,30) are driven by non-vascular factors.

For the other cortical measures in this study, we found no statistically significant effect of menstrual hormone fluctuations, suggesting that, across the menstrual cycle at least, these functions are robust to hormone changes to the limit of our measurement sensitivity. For carotid radius, this replicates previous findings (47), but for OEF, CMRO_2_ and PI, this study represents the first demonstration of stability across the menstrual cycle, possibly reflecting intact cerebrovascular regulation processes.

Regarding the retinal results, conversely to previous work (49), we found no significant interaction between menstrual hormones and vessel density across the retina. However, it should be noted that Guo *et al.,* (49) compared vessel density to menstrual phase rather than hormones, a slightly different comparison that inherently assumes hormone levels. It could be that these observed differences between phases were due to changing hormone levels, rather than the measured hormone level *per se*. For example, Guo *et al.,* (2021) ensured that they tested during the ovulatory phase by monitoring luteinising hormone (LH) levels. The day following the peak was defined as the start of the ovulatory phase. However, the oestrogen peak has been observed to occur, most frequently, two days before ovulation (83). Our results together could therefore indicate that retinal vessel density is sensitive to fluctuating hormone levels (before regulatory mechanisms initiate) rather than the level of hormone itself at the time of testing. This hypothesis could be investigated by utilising high-density sampling across the menstrual cycle, which allows for hormonal changes to be better characterised and has recently been employed in neuroendocrine research (84). Alternatively, and given that OCT-A visualises vascular structure by detecting erythrocyte motion, it would be tempting to speculate that the changes in vessel density reported by Guo et al., (2021) are indeed indirect measurements of a potential change in retinal microvascular flow, in the absence of independent evidence of microvascular remodelling over the course of the menstrual cycle.

Finally, we found that oestrogen significantly decreased retinal blood flow resistance, which suggests that the effect on microvascular perfusion found in the cortex also extends to the retina. A regional analysis found that this result may be driven by the foveal vessels, potentially due to the smaller vessels present. If oestrogen has no impact on larger retinal vessels (or there is a more subtle relationship, as suggested with retinal vessel density), this may overshadow a small effect in the other ROIs. In addition, it is possible that arteries and veins behave differently and since we are not discriminating between them, their effect is balanced out in other ROIs.

As an exploratory aim, we examined whether relationships between the vascular functions differed between low and high hormonal state. We investigated loadings onto PCA components and found a pattern of reduced commonality between the vascular functions in the high endocrine condition compared to the low endocrine condition, possibly suggesting that the inter-function relationships were disrupted in high hormone levels. This may reflect the action of increased hormone levels on some functions more than others, leading to a breakdown in the relationship between those functions. This change in relationship may also suggest a more subtle menstrual cycle-related effect that is not captured by the linear models that use hormone value alone; it may be in response to a change in endocrine state, or a different menstrual-related hormone. However, it is important to note that as an exploratory aim, the current study was not designed to be statistically powered for this type of research question and requires specific investigation in future work.

One of the major motivators for the current study was the potential protective effect of ovarian hormones for neurodegenerative conditions such as dementia and glaucoma (e.g., 1,3,9,10,12,13,85). Multiple studies provide strong evidence that vascular alterations across the eye and brain are associated with dementia or glaucoma risk/diagnosis (86–94). If regularly cycling hormones influence the vascular system, as seen in the current study, and lead to healthier vascular ageing, as suggested by lower cardiovascular and cerebrovascular risk with increased lifetime oestrogen exposure (2,4,8), this may be an avenue in which a neuroprotective effect is delivered.

These results also have implications for other groups that show notable changes in endocrine state, including natural or surgical menopause, pregnancy, puberty, conditions with altered hormone levels (such as polycystic ovary syndrome) and hormone replacement therapy. Many of these states demonstrate greater changes in baseline hormone level than across a regular healthy menstrual cycle, so may also show greater changes in baseline cerebrovascular function. The current study also raises questions about the impact of irregular menstruation, which can be associated with altered hormone levels. Indeed, many of these states and conditions are associated with changes in cardio- and cerebro-vascular disease risk (95–99).

One of the limitations of the current study was that, although the phase days were chosen based on previous literature (e.g., 100), the ovulatory oestrogen peak was missed in multiple participants (see Figure 2), which may have led to an underestimation of oestrogen’s influence. The variation in oestrogen menstrual pattern within and across people has been reported previously (83,101,102) and is an important consideration when using methodologies such as MRI that may afford less scheduling flexibility. This highlights the importance of testing the actual circulating hormone level, rather than making assumptions based on menstrual phase, as this allows direct comparisons to the current hormone level. Additionally, by testing at least three times across a cycle, we were able to capture variation in both oestradiol and progesterone. In order to better investigate these changeable fluctuations in hormones, high-density sampling could be used, in which a smaller number of participants are densely sampled across the cycle to better characterise hormonal fluctuations.

## Conclusion

In conclusion, we found that menstrual-related oestradiol and progesterone had a neuroendocrine influence on multiple baseline vascular functions across brain and eye. This has important implications for menstrual-associated symptoms such as menstrual migraine, as well as cerebrovascular function in other endocrine states. This also suggests a possible mechanism by which these hormones have a neuro- and vaso-protective effect.

## Acknowledgements

We would like to thank Wellcome Trust for their help in publication of this article. This research was funded in whole, or in part, by the Wellcome Trust (WT224267).

MG is supported by the Wellcome Trust [220575/Z/20/Z] and the Engineering and Physical Sciences Research Council [EP/S025901/1].

For the purpose of open access, the author has applied a CC BY public copyright license to any Author Accepted Manuscript version arising from this submission.

## Data statement

Data will be made available upon publication. Code will be available upon request.

## Credit statement

MEW: Conceptualization, Methodology, Investigation, Formal analysis, Writing – original draft, Writing – review & editing

CC: Methodology, Investigation, Formal analysis, Writing – review & editing SD: Project Administration, Investigation, Writing – review & editing

HC: Formal analysis, Writing – review & editing ID: Formal analysis, Writing – review & editing

MG: Methodology, Software, Writing – review & editing YG: Software, Formal analysis, Writing – review & editing DR: Software, Writing – review & editing

MOB: Software, Writing – review & editing LT: Methodology, Writing – review & editing JJS: Conceptualization, Methodology

KM: Conceptualization, Methodology, Formal analysis, Writing – original draft, Writing – review & editing, Supervision, Funding acquisition.

## References

1. Fox M, Berzuini C, Knapp LA. Cumulative estrogen exposure, number of menstrual cycles, and Alzheimer’s risk in a cohort of British women. Psychoneuroendocrinology. 2013 Dec;38(12):2973–82.

2. Muka T, Oliver-Williams C, Kunutsor S, Laven JSE, Fauser BCJM, Chowdhury R, et al. Association of Age at Onset of Menopause and Time Since Onset of Menopause With Cardiovascular Outcomes, Intermediate Vascular Traits, and All-Cause Mortality: A Systematic Review and Meta-analysis. JAMA Cardiol. 2016 Oct 1;1(7):767.

3. Gilsanz P, Lee C, Corrada MM, Kawas CH, Quesenberry CP, Whitmer RA. Reproductive period and risk of dementia in a diverse cohort of health care members. Neurology [Internet]. 2019 Apr 23 [cited 2024 Feb 15];92(17). Available from: https://www.neurology.org/doi/10.1212/WNL.0000000000007326

4. Chen L, Hu Z, Wang X, Zheng C, Cao X, Cai J, et al. Association of age at menarche and menopause, reproductive lifespan, and stroke among Chinese women: Results from a national cohort study [Internet]. Epidemiology; 2023 May [cited 2023 Sep 10]. Available from: http://medrxiv.org/lookup/doi/10.1101/2023.05.23.23290429

5. Schelbaum E, Loughlin L, Jett S, Zhang C, Jang G, Malviya N, et al. Association of Reproductive History With Brain MRI Biomarkers of Dementia Risk in Midlife. Neurology [Internet]. 2021 Dec 7 [cited 2024 Apr 11];97(23). Available from: https://www.neurology.org/doi/10.1212/WNL.0000000000012941

6. Steventon JJ, Lancaster TM, Baker E, Bracher-Smith M, Escott-Price V, Ruth KS, et al. Menopause age, reproductive span and hormone therapy duration predict the volume of medial temporal lobe brain structures in postmenopausal women. Psychoneuroendocrinology. 2023 Sep;106393.

7. 7. Lindseth LRS, De Lange AMG, Van Der Meer D, Agartz I, Westlye LT, Tamnes CK, et al. Female-specific factors are associated with cognition in the UK Biobank cohort [Internet]. 2022 [cited 2024 Apr 11]. Available from: http://medrxiv.org/lookup/doi/10.1101/2022.06.27.22276879

8. Hayward C, Kelly R, Collins P. The roles of gender, the menopause and hormone replacement on cardiovascular function. Cardiovasc Res. 2000 Apr;46(1):28–49.

9. Rocca WA, Bower JH, Maraganore DM, Ahlskog JE, Grossardt BR, De Andrade M, et al. Increased risk of cognitive impairment or dementia in women who underwent oophorectomy before menopause. Neurology. 2007 Sep 11;69(11):1074–83.

10. Rocca WA, Grossardt BR, Shuster LT. Oophorectomy, estrogen, and dementia: A 2014 update. Mol Cell Endocrinol. 2014 May;389(1–2):7–12.

11. El Khoudary SR, Greendale G, Crawford SL, Avis NE, Brooks MM, Thurston RC, et al. The menopause transition and women’s health at midlife: a progress report from the Study of Women’s Health Across the Nation (SWAN). Menopause. 2019 Oct;26(10):1213–27.

12. Pertesi S, Coughlan G, Puthusseryppady V, Morris E, Hornberger M. Menopause, cognition and dementia – A review. Post Reprod Health. 2019 Dec;25(4):200–6.

13. Douglass A, Dattilo M, Feola AJ. Evidence for Menopause as a Sex-Specific Risk Factor for Glaucoma. Cell Mol Neurobiol. 2023 Jan;43(1):79–97.

14. Bassani TB, Bartolomeo CS, Oliveira RB, Ureshino RP. Progestogen-Mediated Neuroprotection in Central Nervous System Disorders. Neuroendocrinology. 2023;113(1):14–35.

15. Okoth K, Smith WP, Thomas GN, Nirantharakumar K, Adderley NJ. The association between menstrual cycle characteristics and cardiometabolic outcomes in later life: a retrospective matched cohort study of 704,743 women from the UK. BMC Med. 2023 Mar 20;21(1):104.

16. Tostes RC, Nigro D, Fortes ZB, Carvalho MHC. Effects of estrogen on the vascular system. Braz J Med Biol Res. 2003 Sep;36(9):1143–58.

17. Robison LS, Gannon OJ, Salinero AE, Zuloaga KL. Contributions of sex to cerebrovascular function and pathology. Brain Res. 2019 May;1710:43–60.

18. Munaut C, Lambert V, Noël A, Frankenne F, Deprez M, Foidart JM, et al. Presence of oestrogen receptor type beta in human retina. Br J Ophthalmol. 2001 Jul;85(7):877–82.

19. Shughrue PJ, Merchenthaler I. Distribution of estrogen receptorimmunoreactivity in the rat central nervous system. J Comp Neurol. 2001 Jul 17;436(1):64–81.

20. Milner TA, Thompson LI, Wang G, Kievits JA, Martin E, Zhou P, et al. Distribution of estrogen receptor beta containing cells in the brains of bacterial artificial chromosome transgenic mice. Brain Res. 2010 Sep;1351:74–96.

21. Mitterling KL, Spencer JL, Dziedzic N, Shenoy S, McCarthy K, Waters EM, et al. Cellular and subcellular localization of estrogen and progestin receptor immunoreactivities in the mouse hippocampus. J Comp Neurol. 2010;NA-NA.

22. Mosconi L, Nerattini M, Matthews DC, Jett S, Andy C, Williams S, et al. In vivo brain estrogen receptor density by neuroendocrine aging and relationships with cognition and symptomatology. Sci Rep. 2024 Jun 20;14(1):12680.

23. Barranca C, Pereira TJ, Edgell H. Oral contraceptive use and menstrual cycle influence acute cerebrovascular response to standing. Auton Neurosci. 2023 Jan;244:103054.

24. Wright ME, Murphy K. A mini-review of the evidence for cerebrovascular changes following gender-affirming hormone replacement therapy and a call for increased focus on cerebrovascular transgender health. Front Hum Neurosci [Internet]. 2023;17. Available from: https://www.frontiersin.org/articles/10.3389/fnhum.2023.1303871

25. Cote S, Fathy K, Lenet S, Espinosa-Bentancourt OE, Lavallee E, Croteau E, et al. OR31-03 Testosterone Gender-affirming Hormonal Therapy Is Associated With Decreased Cerebral Blood Flow In Transgender Teens. J Endocr Soc. 2023 Oct 5;7(Supplement_1):bvad114.2099.

26. Protopopescu X, Butler T, Pan H, Root J, Altemus M, Polanecsky M, et al. Hippocampal structural changes across the menstrual cycle. Hippocampus. 2008 Oct;18(10):985–8.

27. Pletzer B, Kronbichler M, Aichhorn M, Bergmann J, Ladurner G, Kerschbaum HH. Menstrual cycle and hormonal contraceptive use modulate human brain structure. Brain Res. 2010 Aug;1348:55–62.

28. Lisofsky N, Mårtensson J, Eckert A, Lindenberger U, Gallinat J, Kühn S. Hippocampal volume and functional connectivity changes during the female menstrual cycle. NeuroImage. 2015 Sep;118:154–62.

29. Taylor CM, Pritschet L, Olsen RK, Layher E, Santander T, Grafton ST, et al. Progesterone shapes medial temporal lobe volume across the human menstrual cycle. NeuroImage. 2020 Oct;220:117125.

30. Zsido RG, Williams AN, Barth C, Serio B, Kurth L, Mildner T, et al. Ultra-high-field 7T MRI reveals changes in human medial temporal lobe volume in female adults during menstrual cycle. Nat Ment Health. 2023 Oct 5;1(10):761–71.

31. Rizor EJ, Babenko V, Dundon NM, Beverly-Aylwin R, Stump A, Hayes M, et al. Menstrual cycle-driven hormone concentrations co-fluctuate with white and gray matter architecture changes across the whole brain. Hum Brain Mapp. 2024 Aug;45(11):e26785.

32. Franke K, Hagemann G, Schleussner E, Gaser C. Changes of individual BrainAGE during the course of the menstrual cycle. NeuroImage. 2015 Jul;115:1–6.

33. Epperson CN, Haga K, Mason GF, Sellers E, Gueorguieva R, Zhang W, et al. Cortical γ- Aminobutyric Acid Levels Across the Menstrual Cycle in Healthy Women and Those With Premenstrual Dysphoric Disorder: A Proton Magnetic Resonance Spectroscopy Study. Arch Gen Psychiatry. 2002 Sep 1;59(9):851.

34. Harada M, Kubo H, Nose A, Nishitani H, Matsuda T. Measurement of variation in the human cerebral GABA level by in vivo MEGA-editing proton MR spectroscopy using a clinical 3 T instrument and its dependence on brain region and the female menstrual cycle. Hum Brain Mapp. 2011 May;32(5):828–33.

35. De Bondt T, De Belder F, Vanhevel F, Jacquemyn Y, Parizel PM. Prefrontal GABA concentration changes in women—Influence of menstrual cycle phase, hormonal contraceptive use, and correlation with premenstrual symptoms. Brain Res. 2015 Feb;1597:129–38.

36. Zhu X, Wang X, Parkinson C, Cai C, Gao S, Hu P. Brain activation evoked by erotic films varies with different menstrual phases: An fMRI study. Behav BRAIN Res. 2010 Jan;206(2):279–85.

37. Arelin K, Mueller K, Barth C, Rekkas PV, Kratzsch J, Burmann I, et al. Progesterone mediates brain functional connectivity changes during the menstrual cycleâ€”a pilot resting state MRI study. Front Neurosci [Internet]. 2015 Feb 23 [cited 2024 Apr 11];9. Available from: http://journal.frontiersin.org/Article/10.3389/fnins.2015.00044/abstract

38. Sumner RL, McMillan RL, Shaw AD, Singh KD, Sundram F, Muthukumaraswamy SD. Peak visual gamma frequency is modified across the healthy menstrual cycle. Hum BRAIN Mapp. 2018 Aug;39(8):3187–202.

39. Dubol M, Epperson CN, Sacher J, Pletzer B, Derntl B, Lanzenberger R, et al. Neuroimaging the menstrual cycle: A multimodal systematic review. Front Neuroendocrinol. 2021 Jan;60:100878.

40. Haraguchi R, Hoshi H, Ichikawa S, Hanyu M, Nakamura K, Fukasawa K, et al. The Menstrual Cycle Alters Resting-State Cortical Activity: A Magnetoencephalography Study. Front Hum Neurosci. 2021 Jul 26;15.

41. Greenwell S, Faskowitz J, Pritschet L, Santander T, Jacobs EG, Betzel RF. High-amplitude network co-fluctuations linked to variation in hormone concentrations over the menstrual cycle. Netw Neurosci. 2023 Oct 1;7(3):1181–205.

42. Jeong HG, Ham BJ, Yeo HB, Jung IK, Joe SH. Gray matter abnormalities in patients with premenstrual dysphoric disorder: An optimized voxel-based morphometry. J Affect Disord. 2012 Nov;140(3):260–7.

43. Liu P, Wei Y, Fan Y, Liao H, Wang G, Li R, et al. Cortical and subcortical changes in patients with premenstrual syndrome. J Affect Disord. 2018 Aug;235:191–7.

44. 44. Marjanovic SB, Bukhari MCH, Kjelkenes R, Voldsbekk I, Barth C, Westlye LT. Assessing brain morphological correlates of premenstrual symptoms in young healthy females [Internet]. 2024 [cited 2024 Nov 29]. Available from: https://osf.io/h9w4v

45. Brackley KJ, Ramsay MM, Broughton Pipkin F, Rubin PC. The effect of the menstrual cycle on human cerebral blood flow: studies using Doppler ultrasound. Ultrasound Obstet Gynecol. 1999 Jul;14(1):52–7.

46. Otomo M, Harada M, Abe T, Matsumoto Y, Abe Y, Kanazawa Y, et al. Reproducibility and Variability of Quantitative Cerebral Blood Flow Measured by Multi-delay 3D Arterial Spin Labeling According to Sex and Menstrual Cycle. J Med Invest. 2020;67(3.4):321–7.

47. Cote S, Butler R, Michaud V, Lavallee E, Croteau E, Mendrek A, et al. The regional effect of serum hormone levels on cerebral blood flow in healthy nonpregnant women. Hum Brain Mapp. 2021 Dec;42(17):5677–88.

48. Wagner SK, Fu DJ, Faes L, Liu X, Huemer J, Khalid H, et al. Insights into Systemic Disease through Retinal Imaging-Based Oculomics. Transl Vis Sci Technol. 2020 Feb 12;9(2):6.

49. Guo L, Zhu C, Wang Z, Gao Z, Zhang Z, Pan Q. Retinal Vascular Changes during the Menstrual Cycle Detected with Optical Coherence Tomography Angiography. Kusuhara S, editor. J Ophthalmol. 2021 Jul 12;2021:1–8.

50. Giarratano Y, Bianchi E, Gray C, Morris A, MacGillivray T, Dhillon B, et al. Automated Segmentation of Optical Coherence Tomography Angiography Images: Benchmark Data and Clinically Relevant Metrics. Transl Vis Sci Technol. 2020 Dec 3;9(13):5.

51. Giarratano Y, Pavel A, Lian J, Andreeva R, Fontanella A, Sarkar R, et al. A Framework for the Discovery of Retinal Biomarkers in Optical Coherence Tomography Angiography (OCTA). In: Fu H, Garvin MK, MacGillivray T, Xu Y, Zheng Y, editors. Ophthalmic Medical Image Analysis. Cham: Springer International Publishing; 2020. p. 155–64.

52. Gruber CJ, Tschugguel W, Schneeberger C, Huber JC. Production and Actions of Estrogens. N Engl J Med. 2002 Jan 31;346(5):340–52.

53. Joris P, Mensink R, Adam T, Liu T. Cerebral Blood Flow Measurements in Adults: A Review on the Effects of Dietary Factors and Exercise. Nutrients. 2018 Apr 25;10(5):530.

54. Lu H, Ge Y. Quantitative evaluation of oxygenation in venous vessels using T2- Relaxation-Under-Spin-Tagging MRI. Magn Reson Med. 2008 Aug;60(2):357–63.

55. Lu H, Xu F, Grgac K, Liu P, Qin Q, Van Zijl P. Calibration and validation of TRUST MRI for the estimation of cerebral blood oxygenation. Magn Reson Med. 2012 Jan;67(1):42–9.

56. Xu F, Li W, Liu P, Hua J, Strouse JJ, Pekar JJ, et al. Accounting for the role of hematocrit in between-subject variations of MRI-derived baseline cerebral hemodynamic parameters and functional BOLD responses. Hum Brain Mapp. 2018 Jan;39(1):344–53.

57. Whittaker JR, Fasano F, Venzi M, Liebig P, Gallichan D, Möller HE, et al. Measuring Arterial Pulsatility With Dynamic Inflow Magnitude Contrast. Front Neurosci. 2022 Jan 17;15:795749.

58. Bianciardi M, Toschi N, Polimeni JR, Evans KC, Bhat H, Keil B, et al. The pulsatility volume index: an indicator of cerebrovascular compliance based on fast magnetic resonance imaging of cardiac and respiratory pulsatility. Philos Trans R Soc Math Phys Eng Sci. 2016 May 13;374(2067):20150184.

59. Cox RW. AFNI: Software for Analysis and Visualization of Functional Magnetic Resonance Neuroimages. Comput Biomed Res. 1996 Jun;29(3):162–73.

60. Pinto J, Chappell MA, Okell TW, Mezue M, Segerdahl AR, Tracey I, et al. Calibration of arterial spin labeling data—potential pitfalls in post-processing. Magn Reson Med. 2020 Apr;83(4):1222–34.

61. Herscovitch P, Raichle ME. What is the Correct Value for the Brain-Blood Partition Coefficient for Water? J Cereb Blood Flow Metab. 1985 Mar;5(1):65–9.

62. Zhao JM, Clingman CS, Närväinen MJ, Kauppinen RA, Van Zijl PCM. Oxygenation and hematocrit dependence of transverse relaxation rates of blood at 3T. Magn Reson Med. 2007 Sep;58(3):592–7.

63. Buxton RB, Frank LR, Wong EC, Siewert B, Warach S, Edelman RR. A general kinetic model for quantitative perfusion imaging with arterial spin labeling. Magn Reson Med. 1998 Sep;40(3):383–96.

64. Jenkinson M, Beckmann CF, Behrens TEJ, Woolrich MW, Smith SM. FSL. NeuroImage. 2012 Aug;62(2):782–90.

65. Desikan RS, Ségonne F, Fischl B, Quinn BT, Dickerson BC, Blacker D, et al. An automated labeling system for subdividing the human cerebral cortex on MRI scans into gyral based regions of interest. NeuroImage. 2006 Jul;31(3):968–80.

66. Izzo R, Steinman D, Manini S, Antiga L. The Vascular Modeling Toolkit: A Python Library for the Analysis of Tubular Structures in Medical Images. J Open Source Softw. 2018 May 26;3(25):745.

67. Li A, You J, Du C, Pan Y. Automated segmentation and quantification of OCT angiography for tracking angiogenesis progression. Biomed Opt Express. 2017 Dec 1;8(12):5604.

68. R core team. R: A language and environment for statistical computing. [Internet]. Vienna, Austria.: R Foundation for Statistical Computing; 2022. Available from: https://www.R-project.org/

69. Kuznetsova A, Brockhoff PB, Christensen RHB. **lmerTest** Package: Tests in Linear Mixed Effects Models. J Stat Softw [Internet]. 2017 [cited 2023 Oct 9];82(13). Available from: http://www.jstatsoft.org/v82/i13/

70. Lenton EA, Sulaiman R, Sobowale O, Cooke I. The human menstrual cycle: plasma concentrations of prolactin, LH, FSH, oestradiol and progesterone in conceiving and non- conceiving women. Reproduction. 1982;65(1):131–9.

71. Lee SJ, Lenton EA, Sexton L, Cooke ID. The effect of age on the cyclical patterns of plasma LH, FSH, oestradiol and progesterone in women with regular menstrual cycles. Hum Reprod. 1988 Oct;3(7):851–5.

72. Chi OZ, Barsoum S, Wen Y, Liu X, Weiss HR. 17beta-estradiol prevents blood-brain barrier disruption induced by VEGF. Horm Metab Res Horm Stoffwechselforschung Horm Metab. 2004 May;36(5):272–6.

73. Hurn PD, Littleton-Kearney MT, Kirsch JR, Dharmarajan AM, Traystman RJ. Postischemic Cerebral Blood Flow Recovery in the Female: Effect of 17β-Estradiol. J Cereb Blood Flow Metab. 1995 Jul;15(4):666–72.

74. Santizo RA, Xu HL, Galea E, Muyskens S, Baughman VL, Pelligrino DA. Combined Endothelial Nitric Oxide Synthase Upregulation and Caveolin-1 Downregulation Decrease Leukocyte Adhesion in Pial Venules of Ovariectomized Female Rats. Stroke. 2002 Feb;33(2):613–6.

75. Chen Z, Xi G, Mao Y, Keep RF, Hua Y. Effects of Progesterone and Testosterone on ICH-Induced Brain Injury in Rats. In: Zhang J, Colohan A, editors. Intracerebral Hemorrhage Research [Internet]. Vienna: Springer Vienna; 2011 [cited 2023 Sep 11]. p. 289–93. (Acta Neurochirurgica Supplementum; vol. 111). Available from: https://link.springer.com/10.1007/978-3-7091-0693-8_48

76. Chen Y, Herrold AA, Gallagher V, Martinovich Z, Bari S, Vike NL, et al. Preliminary Report: Localized Cerebral Blood Flow Mediates the Relationship between Progesterone and Perceived Stress Symptoms among Female Collegiate Club Athletes after Mild Traumatic Brain Injury. J Neurotrauma. 2021 Jul 1;38(13):1809–20.

77. Gibson CL, Coomber B, Murphy SP. Progesterone is neuroprotective following cerebral ischaemia in reproductively ageing female mice. Brain. 2011 Jul 1;134(7):2125–33.

78. Cunha TRD, Giesen JAS, Rouver WN, Costa ED, Grando MD, Lemos VS, et al. Effects of progesterone treatment on endothelium-dependent coronary relaxation in ovariectomized rats. Life Sci. 2020 Apr;247:117391.

79. Da Costa DT, Gonçalves LT, Giesen JAS, Dos Santos RL. Progesterone modulates endothelium-dependent coronary vascular reactivity in SHR. J Mol Endocrinol. 2021 Feb;66(2):171–80.

80. Gibson CL, Constantin D, Prior MJW, Bath PMW, Murphy SP. Progesterone suppresses the inflammatory response and nitric oxide synthase-2 expression following cerebral ischemia. Exp Neurol. 2005 Jun;193(2):522–30.

81. Sunday L, Tran MM, Krause DN, Duckles SP. Estrogen and progestagens differentially modulate vascular proinflammatory factors. Am J Physiol-Endocrinol Metab. 2006 Aug;291(2):E261–7.

82. Davis TL, Kwong KK, Weisskoff RM, Rosen BR. Calibrated functional MRI: Mapping the dynamics of oxidative metabolism. Proc Natl Acad Sci. 1998 Feb 17;95(4):1834–9.

83. Maman E, Adashi EY, Baum M, Hourvitz A. Prediction of ovulation: new insight into an old challenge. Sci Rep. 2023 Nov 15;13(1):20003.

84. Pritschet L, Taylor CM, Santander T, Jacobs EG. Applying dense-sampling methods to reveal dynamic endocrine modulation of the nervous system. Curr Opin Behav Sci. 2021 Aug;40:72–8.

85. Fotesko K, Thomsen BSV, Kolko M, Vohra R. Girl Power in Glaucoma: The Role of Estrogen in Primary Open Angle Glaucoma. Cell Mol Neurobiol [Internet]. 2020 Oct 11 [cited 2021 Jan 15]; Available from: http://link.springer.com/10.1007/s10571-020-00965-5

86. Piltz-seymour JR, Grunwald JE, Hariprasad SM, Dupont J. Optic nerve blood flow is diminished in eyes of primary open-angle glaucoma suspects. Am J Ophthalmol. 2001 Jul;132(1):63–9.

87. Shiga Y, Omodaka K, Kunikata H, Ryu M, Yokoyama Y, Tsuda S, et al. Waveform Analysis of Ocular Blood Flow and the Early Detection of Normal Tension Glaucoma. Investig Opthalmology Vis Sci. 2013 Nov 21;54(12):7699.

88. Yamashita K ichiro, Taniwaki Y, Utsunomiya H, Taniwaki T. Cerebral Blood Flow Reduction Associated with Orientation for Time in Amnesic Mild Cognitive Impairment and Alzheimer Disease Patients: Disorientation Related Hypoperfusion Analyzed by 3D- SSP. J Neuroimaging. 2014 Nov;24(6):590–4.

89. 89. Nakazawa T. Ocular Blood Flow and Influencing Factors for Glaucoma: Asia-Pac J Ophthalmol. 2016;5(1):38–44.

90. Chandler HL, Wise RG, Murphy K, Tansey KE, Linden DEJ, Lancaster TM. Polygenic impact of common genetic risk loci for Alzheimer’s disease on cerebral blood flow in young individuals. Sci Rep. 2019 Dec;9(1):467.

91. 91. Chandler HL, Wise RG, Linden DE, Williams J, Murphy K, Lancaster TM, et al. Alzheimer’s genetic risk effects on cerebral blood flow are spatially consistent and proximal to gene expression across the lifespan [Internet]. 2021 [cited 2024 Sep 2]. Available from: http://biorxiv.org/lookup/doi/10.1101/2020.12.31.424949

92. Ortner M, Hauser C, Schmaderer C, Muggenthaler C, Hapfelmeier A, Sorg C, et al. Decreased Vascular Pulsatility in Alzheimer’s Disease Dementia Measured by Transcranial Color-Coded Duplex Sonography. Neuropsychiatr Dis Treat. 2019 Dec;Volume 15:3487–99.

93. Wang Q, Qu X, Wang H, Chen W, Sun Y, Li T, et al. Arterial spin labeling reveals disordered cerebral perfusion and cerebral blood flow-based functional connectivity in primary open-angle glaucoma. Brain Imaging Behav [Internet]. 2023 Nov 25 [cited 2023 Dec 2]; Available from: https://link.springer.com/10.1007/s11682-023-00813-2

94. Desai B, Edwards O, Beishon L. The role of cerebral blood flow in the pathogenesis of Alzheimer’s Disease Dementia. Aging Health Res. 2024 Jun;4(2):100188.

95. Connelly P, Freel E, Perry C, Ewan J, Touyz R, Currie G, et al. Gender-Affirming Hormone Therapy, Vascular Health and Cardiovascular Disease in Transgender Adults. HYPERTENSION. 2019 Dec;74(6):1266–74.

96. Zhang J, Xu J, Qu Q, Zhong G. Risk of Cardiovascular and Cerebrovascular Events in Polycystic Ovarian Syndrome Women: A Meta-Analysis of Cohort Studies. Front Cardiovasc Med. 2020 Nov 12;7.

97. Kilic D, Kilic ID, Sevgican CI, Kilic O, Alatas E, Arslan M, et al. Arterial stiffness measured by cardio-ankle vascular index is greater in non-obese young women with polycystic ovarian syndrome. J Obstet Gynaecol Res. 2021 Feb;47(2):521–8.

98. Pribish AM, Iwamoto SJ. Cardiovascular disease and feminizing gender-affirming hormone therapy: implications for the provision of safe and lifesaving care. Curr Opin Physiol. 2023 Jun;33:100650.

99. Wright ME, Richards CT, Davies S, Lord RN, Rees A, Murphy K. Cortical vasculature in polycystic ovary syndrome and associations with circulating testosterone. Endocr Abstr [Internet]. 2025 Feb 19 [cited 2025 Feb 26]; Available from: http://www.endocrine-abstracts.org/ea/0109/ea0109P197

100. Draper CF, Duisters K, Weger B, Chakrabarti A, Harms AC, Brennan L, et al. Menstrual cycle rhythmicity: metabolic patterns in healthy women. Sci Rep. 2018 Dec;8(1):14568.

101. Gandara BK, Leresche L, Mancl L. Patterns of Salivary Estradiol and Progesterone across the Menstrual Cycle. Ann N Y Acad Sci. 2007 Mar;1098(1):446–50.

102. Celec P, Ostaniková D, Skoknová M, Hodosy J, Putz Z, Kúdela M. Salivary Sex Hormones during the Menstrual Cycle. Endocr J. 2009;56(3):521–3.

103. Smith SM, Jenkinson M, Woolrich MW, Beckmann CF, Behrens TEJ, Johansen-Berg H, et al. Advances in functional and structural MR image analysis and implementation as FSL. NeuroImage. 2004 Jan;23:S208–19.

104. Woolrich MW, Jbabdi S, Patenaude B, Chappell M, Makni S, Behrens T, et al. Bayesian analysis of neuroimaging data in FSL. NeuroImage. 2009 Mar;45(1):S173–86.

105. Zhang Y, Brady M, Smith S. Segmentation of brain MR images through a hidden Markov random field model and the expectation-maximization algorithm. IEEE Trans Med Imaging. 2001 Jan;20(1):45–57.

