## Supplemental material for "Endocrine modulation of cortical and retinal blood flow across the menstrual cycle"

### Methods

*Grey matter mask generation*

The T1 MPRAGE structural images were processed using the fsl_anat pipeline (64,103,104), which includes reorientating to standard orientation, bias-field correction, brain extraction, and tissue-type segmentation (FAST; 105). This generated a grey matter mask to be used to mask cortical atlas structures and grey matter CBF.

### Results

*Haemoglobin levels*

It was separately investigated whether the estimated Hb values used within the OEF and $CMRO_2$ calculation were independently influenced by hormone level and could be biassing the results. The influence of oestradiol and resProgesterone (random effects) on Hb levels was investigated using linear mixed models (65,66), with participant as a fixed effect. Neither oestradiol ($\chi^2(1)$= 1.241; p=0.265) nor resProgesterone ($\chi^2(1)$= 1.995; p=0.158) significantly contributed to haematocrit variance, suggesting that this was not biasing the OEF and $CMRO_2$ results.

*End-tidal $CO_2$*

Baseline partial pressures of end-tidal $CO_2$ traces were measured from expirations collected using a facemask and an AD Instruments gas analyser and data sampling system (PowerLab®, ADInstruments, Sydney, Australia). This was collected for 20 participants, with 49 total datapoints. In order to examine whether endocrine changes in blood $CO_2$ could be driving our CBF result, 500 seconds of baseline recording were averaged over (median) and linear mixed models used to investigate whether oestradiol or resProgesterone explained a significant amount of variance (using participant as a fixed effect). It was found that neither oestradiol ($\chi^2(1)$= 1.409; p=0.235) nor resProgesterone ($\chi^2(1)$= 1.186; p=0.276) significantly contributed to end-tidal $CO_2$ variance.

*Exploratory relationships analysis*

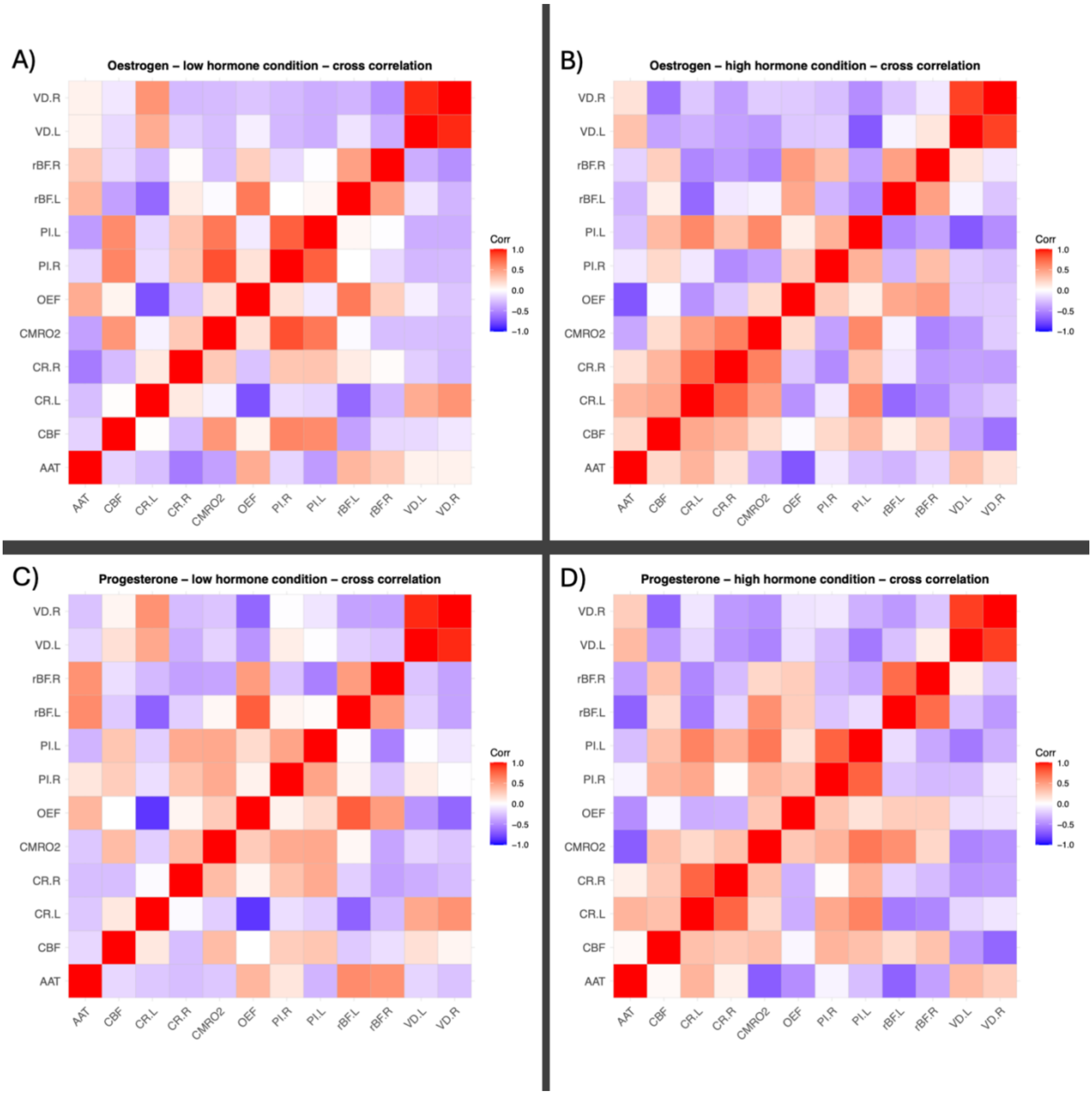


*Figure S1* – A cross-correlation matrix of each condition (low oestrogen, high oestrogen, low progesterone, high progesterone) to be analysed in the principal component analysis. The correlation statistic in a Person's rho, with more red values indicated more positive correlations, and more blue values indicating more negative correlations. AAT=Arterial arrival time; CBF=Cerebral blood flow; CR.L=Carotid radius left; CR.R=Carotid radius right; $CMRO_2$=Cerebral metabolic rate of oxygen; OEF=Oxygen extraction fraction; PI.L=Pulsatility index left; PI.R=Pulsatility index right; rBF.L=Retinal blood flow resistance left; rBF.R=Retinal blood flow resistance right; VD.L=Retinal vessel density left; VD.L=Retinal vessel density left.

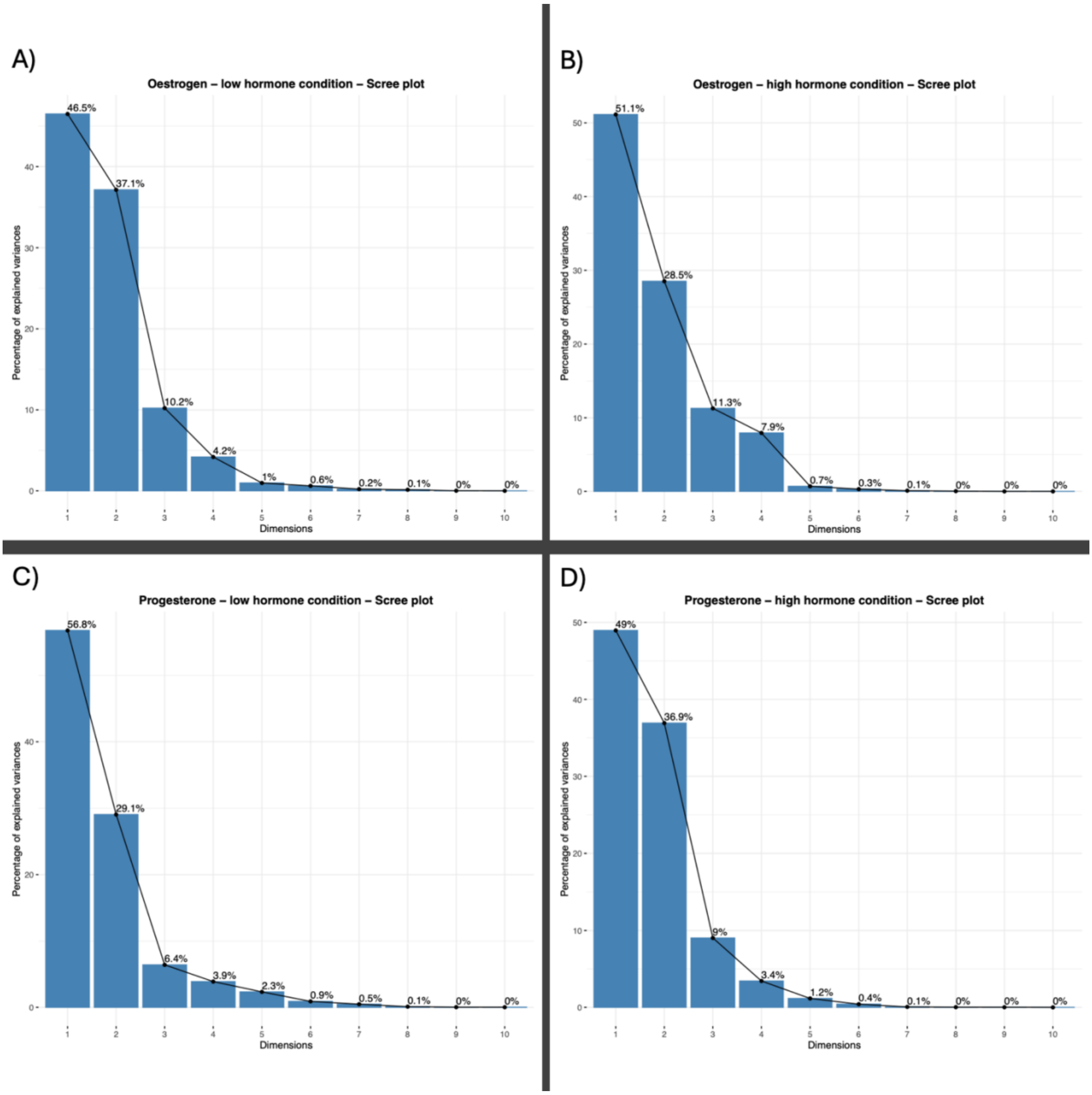


*Figure S2* – Scree plots for each of the four conditions (low oestrogen, high oestrogen, low progesterone, high progesterone) following principal component analysis (PCA), illustrating the amount of variance explained by the generated dimensions/components.
